# Enhancing Variant Calling in Whole Exome Sequencing (WES) Data Using Population-Matched Reference Genomes

**DOI:** 10.1101/2024.08.19.608554

**Authors:** Shuming Guo, Zhuo Huang, Yanming Zhang, Yukun He, Xiangju Chen, Wenjuan Wang, Lansheng Li, Yu Kang, Zhancheng Gao, Jun Yu, Zhenglin Du, Yanan Chu

## Abstract

Whole exon sequencing (WES) data are frequently used for cancer diagnosis and genome-wide association studies (GWAS), hinging upon high-coverage read mapping, informative variant calling, and high-quality reference genomes. The center position of the currently used genome assembly, GRCh38, is now challenged by two newly publicized telomere-to-telomere or T2T genomes, T2T-CHM13 and T2T-YAO, and it becomes urgent to have a comparative study to test population specificity using the three reference genomes based on real case WES data. We here report our analysis along this line for 19 tumor samples collected from Chinese patients. The primary comparison of the exon regions among the three references reveals that the sequences in up to ∼1% target regions in YAO are widely diversified from GRCh38 and may lead to off-target in sequence capture. However, YAO still outperforms GRCh38 genomes by obtaining 7.41% more mapped reads. Due to more reliable read-mapping and closer phylogenetic relationship with the samples than GRCh38, YAO reduces half of variant calls of clinical significance which are mostly benign while keeping sensitivity in identifying pathogenic variants. YAO also outperforms CHM13 in reducing calls of Chinese-specific variants. Our findings highlight the critical need for employing population-specific reference genomes in genomic analysis to ensure accurate variant analysis and the significant benefits of tailoring these approaches to the unique genetic backgrounds of each ethnic group.

## Introduction

Next-generation sequencing (NGS) has been extensively employed in a wide spectrum of clinical applications [1, 2]. More and more practice of precision medicine, including diagnosis, prognosis, and therapy selection across genetic disorders, oncology, and infectious diseases relies on sequencing of human genome DNA [3, 4]. Both whole genome sequencing (WGS) and whole exome sequencing (WES) are widely used to identify genetic (germline) or somatic (such as in tumor tissues) variations in helping genetic disorder diagnosis or discovering novel tumor antigens [2, 5–7]. The WES, which only sequences the protein-coding region (about 1-2 % of the whole human genome) by targeting enrichment, costs much less and is more widely applied clinically [8].

For human and other animals with large genomes, analyses of high-throughput data start with mapping sequencing reads against a reference genome, which is the foundation for all resequencing data analyses for biomedical research and clinical applications. As such, pursuing a complete and accurate human genome reference has been a long-lasting goal for the society of biomedicine. The Genome Reference Consortium (GRC) has continuously improved the human reference genome from the first version by Human Genome Project in 2001 to the up-to-date GRCh38 released in 2013 [9–11]. In 2022, advents the first complete human genome haplotype—CHM13 which is a telomere-to-telomere (T2T) assembly of the European ancestry genome from a hydatidiform mole-CHM13 with unprecedented high-quality of Q73.94, i.e., it has less than one error in 24.8-Mbp sequence [12]. Next year, the complete sequence of chromosome Y from the HG002 genome (European Jewish ancestry) is added to cover all chromosomes (22+XY) of human, leading to T2T-CHM13 v2.0 [13]; independently, our group completed the assembly of the diploid human genome T2T-YAO, based on data from a trio from Han Chinese ancestry, and achieved a comparable high quality of a haplotype version— YAO-hp (Q74.69, i.e., one error in 29.5-Mbp sequence) [14]. And more efforts have been made to create reference genome for the Han population, including Han1 [15] and CN1 [16], with lower quality. Furthermore, a draft human pangenome reference [17] and a comprehensive pangenome reference encompassing 36 Chinese populations have been developed [18], providing valuable resources for understanding genetic diversity across different populations.

It is presumed that the high quality and completeness of human reference genome will improve the accuracy in read-mapping and variant-calling for the analysis of high throughput sequencing data [19]. A reference genome of closer phylogenetic relationship will theoretically abate the number of unmapped reads and improve mapping quality by reducing ambiguous mapping of reads with mismatches. Given the great degree of global genetic variation, reference genomes representative of populations is necessary for effectively performing omics analyses on those populations [14, 20–23]. CHM13 has been publicized as a major improvement from the currently used GRCh38, while YAO is closer to Chinese and of comparable quality to CHM13, potentiating improvements in genomic analysis for Chinese by substituting the current GRCh38 reference. However, the improvement in using higher-quality reference genome with closer phylogenetic relationship has not yet been quantified especially for samples from Chinese.

To evaluate the improvement provided by new reference genomes, we designed a study to quantitatively assess the differences among three genomes when analyzing Whole Exome Sequencing (WES) data from Chinese samples. We selected WES over Whole Genome Sequencing (WGS) because WES, or targeted sequencing of gene panels, is the most prevalent practice in clinical personalized medicine. The impact of different reference genomes on this specific application, as well as the bias introduced by capture probes designed with GRCh38 for WES or panel sequencing in Chinese populations, remains largely unexplored. Previous studies have investigated the performance of various references using standard benchmark genomes, such as HG002 and HG005, as well as WGS data from public population datasets [16, 19], leading us to avoid redundant analyses. In this study, we use the complete human haplotypes of T2T-YAO-hp, T2T-CHM13 v2.0, and GRCh38 (excluding decoy genome), all of which include a single copy of 22+XY chromosomes. For brevity, these references are referred to as YAO, CHM13, and GRCh38. We first compared the basic statistics of these references, particularly their coverage of exomes. We then utilized a WES dataset from 19 gastric tumor samples from Han Chinese, aligning the data separately against each of the three references for initial evaluation.

The reliance on GRCh38 of current standard variant calling processes, despite extensively optimized and evaluated, warrants reassessment when using alternative references. Consequently, we compared the performance of three reference genomes in parallel, analyzing each step of the variant calling process—from mapping to raw variants and final variants after filtering with default cutoffs. Variants in homozygous, heterozygous, and somatic categories were compared both in whole genome (target and flanking regions) and only in target regions. Significant differences were observed across all comparison matrices when using different references. Although this study did not achieve an optimized procedure for WES analysis using alternative reference to GRCh38, our results highlight the urgent need for establishing population-specific reference genomes for Chinese populations.

## Results and Discussion

### Basic statistics of YAO in comparison to GRCh38 and CHM13

The lengths of the three genome assemblies are as follows: the longest is 3,117,292,070 bp for CHM13, followed by 3,088,286,401 bp for GRCh38 (including 150,630,719 Ns), and the shortest is 3,062,724,542 bp for YAO (**Table S1**). Among these references, YAO is derived from a real individual, whereas CHM13 and GRCh38 are not, with differences in length of less than 2%. Variability in chromosome length is well-documented and is primarily attributed to the expansion and contraction of highly repetitive regions, particularly in centromeric and heterochromatic areas. Notable examples include the megabase-long expansion on chromosome 9 in CHM13 [12] and the extensive length diversity observed on chromosome Y [24]. The GC content, which represents the fraction of guanine (G) and cytosine (C) nucleotides, varies across different regions of the human genome and plays a significant role in the efficiency of Illumina sequencing and subsequent analysis. YAO and CHM13 have similar GC content, 40.75% and 40.79%, respectively, slightly lower than that of GRCh38 (41.59%), possibly due to the fully-filled sequences of the relatively AT-rich centromere regions in the two better-assembled genomes. Based on the up-to-date annotation files [25–27], the collective exon lengths (including exons of both protein-coding and non-coding genes) of the three genomes are the longest GRCh38, 156,332,309 bp (5.062% of the genome length), the next YAO, 156,053,407 bp (5.095% of the genome length), and the shortest CHM13, 153,061,925 bp (4.910% of the genome length).

To facilitate the subsequent comparison of WES dataset analysis across the three references, we focused on the exon regions of protein-coding genes targeted by Agilent kit of SureSelect Human All Exon V6 and lifted their original coordinates in GRCh37 to the three reference genomes (**Table S2**). Of the 243,190 targeting regions in a collective length of 60,700,153 bp in GRCh37, 99% can be successfully lifted to all three references. There are only 1,700 uncertain regions in GRCh37, either mapped to multiple sites or unmappable to CHM13 and YAO, which are more than the unmappable 1,281 regions in GRCh38 (**Table S2**). Nevertheless, all three reference genomes retain >60Mb total targetable exon sequences, and the difference among them is rather relatively neglectable. We also calculated sequence identity for each lifted region as YAO *vs*. CHM13 and YAO *vs*. GRCh38. Although ∼85% of the lifted regions are strictly conserved to show 100% identity, there are ∼0.6% regions with sequence identity <80% (1,602 regions to GRCh38 and 1437 regions to CHM13, **Fig. 1**). Together with the 1,705 failed regions, 1-2% of the targetable regions where the capture probes are designed according to the GRCh37/38 genome do not match the samples from Chinese individuals, suggesting potential underrepresentation in their WES data of these regions.

**Fig 1.**
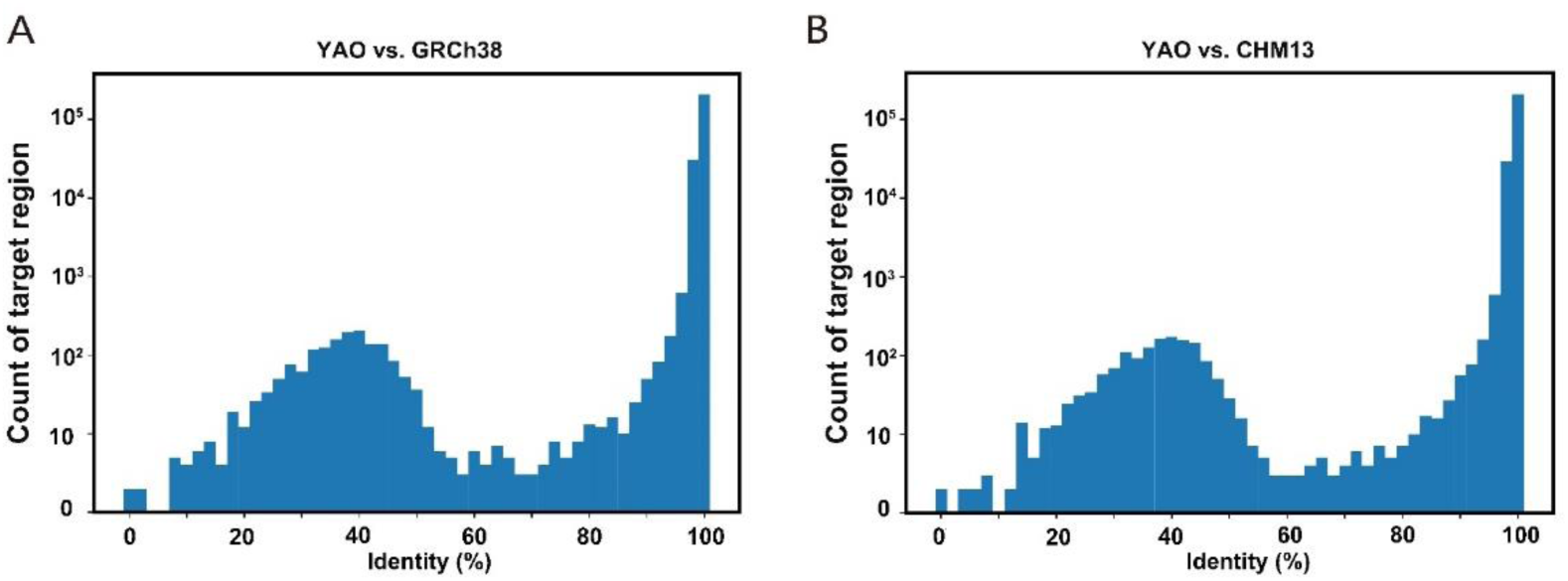
Sequence identity among WES target regions of YAO, CHM13, and GRCh38. **A.** YAO vs. GRCh38. **B.** YAO vs. CHM13. The coordinate information of target regions from the Agilent SureSelect Human All Exon V6 was lifted from GRCh37 to YAO, CHM13, and GRCh38 reference genomes using the transanno tool.

### WES data and alignment to the references

A collection of 19 paraffin embedded gastric tumor samples, 9 benign gastric stromal tumors and 10 malignant gastric cancers, from Han Chinese patients in Linfen Central Hospital were applied to DNBSEQ-T7 platform for 150 bp pair-end WES sequencing (**Table S3**). Data analysis followed the process shown in **Fig S1**. The quality of sequencing reads has an average of 94.8% of Phred value >Q30 and the average sequencing yield is 17.7 ± 5.05 Gb after trimming off bases below Q20, equal to ∼ 300 × sequencing depth of the target regions. The difference between the two groups is not significant in both sequencing yield (19.45 ± 2.06 Gb *vs.* 18.69 ± 5.14 Gb, *p*=0.691, t-test) and quality (Q33.3 ± 0.68 vs. Q33.7 ± 0.71, *p*=0.815, t-test).

The first step in NGS data analysis is aligning the sequencing reads against a reference genome. It is well known that a small percentage of the sequencing reads cannot be mapped to the human reference genome in practical analysis due to incompleteness and misassembling of reference, and it has been suggested that improving the human reference genome may also improve the alignment rate [19]. We first mapped the clean sequencing data separately to YAO, CHM13, and GRCh38 and compared their mapping and mismatch rates. On average, a total of 17.87 ± 3.89 Gb bases are mapped to YAO, which is 5.3 Mb on average more than that to GRCh38 (p=5.945×10^-5^, paired t-test) and almost identical to CHM13 (p=0.093, paired t-test) (**Fig. S2A**). In addition, the average mismatch rate (mismatched bases in aligned reads / total aligned bases) of reads alignment against YAO is 0.214 ± 0.013%, showing significant improvement when compared to GRCh38 (0.245± 0.016%, p=2.79×10^-15^, paired t-test) and CHM13 (0.227± 0.013%, p=1.65×10^-23^, paired t-test) (**Fig.S2B**). Although the differences are subtle but very significant as each sample shows reduced number of mismatches when aligned to YAO vs. to CHM13 and GRCh38.

Greater improvement in mapping becomes more obvious after the removal of low-quality reads (MAPQ <20). On average, the mapped bases are 17.9 ± 3.89 Gbp when aligned against YAO, resulting in 3.37Mbp and 1.23Gb additional aligned bases against CHM13 (p=8.95×10^-3^, paired t-test) and GRCh38 (p=1.33×10^-13^, paired t-test), equal to 0.02% and 7.41% improvements, respectively (**Fig. 2A**). The average mismatch rate in the high-quality mapped reads against YAO is reduced to 0.204 ± 0.0142%, significantly lower than that of 0.215 ± 0.0141% (p=2.01×10^-21^, paired t-test) against CHM13 and 0.222 ± 0.0150% (p=3.27×10^-18^, paired t-test) against GRCh38 (**Fig. 2B**).

**Fig 2.**
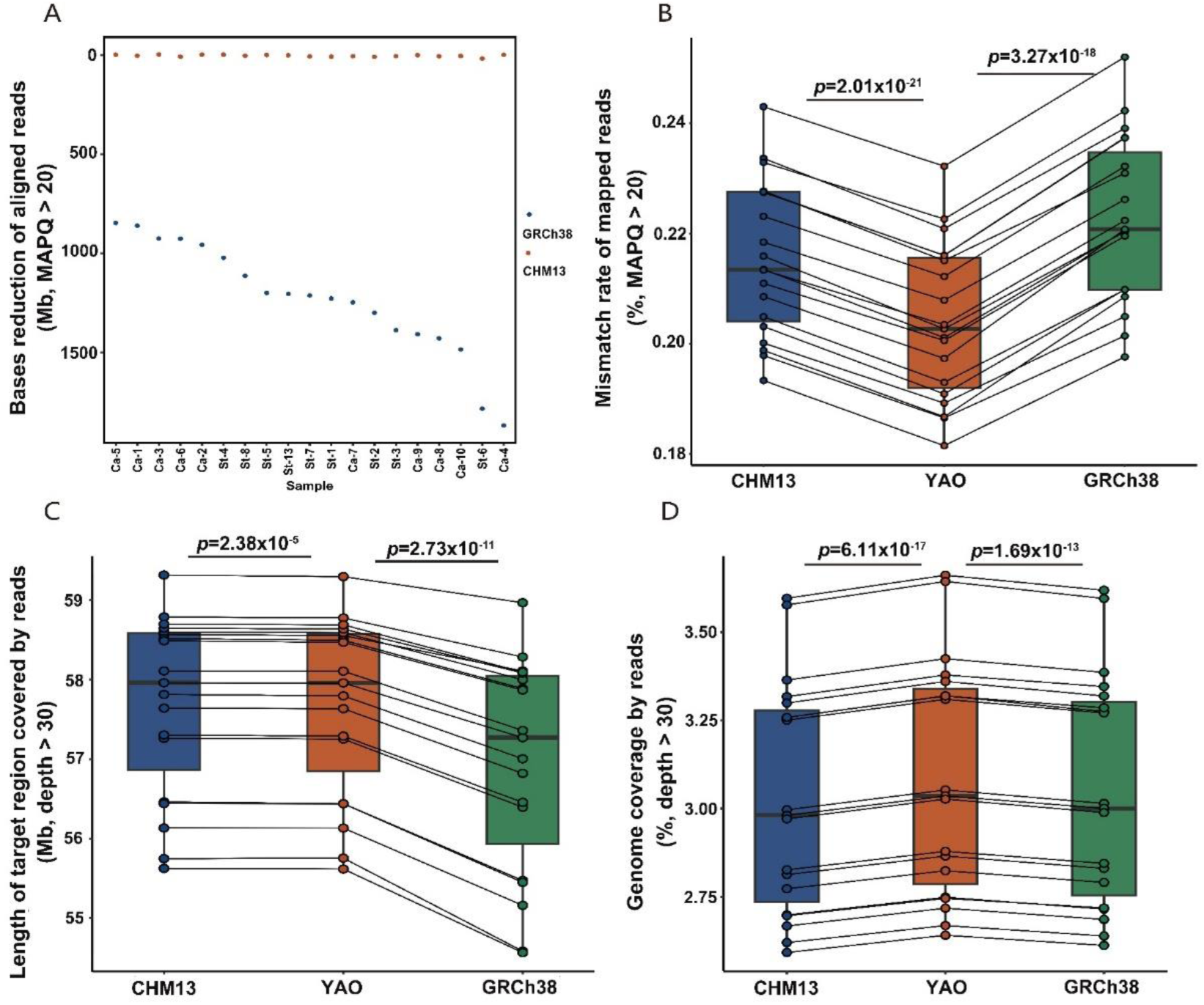
Comparison of read alignment referencing CHM13, YAO, and GRCh38. **A**. The reduction of base of aligned reads (MAPQ>20) in each sample referencing CHM13 and GRCh38 when compared to YAO in whole genome region. Blue and orange solid circles represent the difference when comparing YAO vs. GRCh38 and YAO vs. CHM13, respectively. Samples are sorted according to their total mapped reads. St, gastric stromal tumor; Ca, gastric cancer. **B.** Mismatch rates of aligned reads (MAPQ>20) across the three reference genomes, calculated as the number of mismatched bases divided by the total number of aligned bases. **C.** The length of exon regions with sequencing depth > 30 in target regions. **D.** The fraction of total genomic regions covered by reads with depth > 30 in the three reference genomes. The *p* value of paired T-test is labeled above each comparison. The points representing the same individual across different reference genome are connected by solid lines

Focusing on the target exon region lifted from GRCh37 to CHM13, YAO and GRCh38, we found 1-5 Mbp regions in each sample failed to be sufficiently covered by the reads (depth < 30×), regardless of the reference genome used. This confirms the existence of off-target effect in the capture process of target sequencing due to unmatched probes against the Chinese samples (**Fig. 2C**). In addition to the target exon sequences, for which the capture probes are designed, WES reads often cover the flanking area due to the *hitchhiker* DNA fragments captured by the probes. Despite not being fully targeted in the enrichment process due to unmatched probes, WES reads from the 19 Chinese samples still cover 45.86±7.49% genomic regions in YAO, significantly longer than those in CHM13 and GRCh38 (**Fig. S2C**). After the exclusion of total regions with a sequencing depth less than 30× for calling reliable variants, 3.09±0.33% genome length of YAO remains covered, which is significantly longer than those of CHM13 (3.03±0.32%, p=6.11×10^-17^) and GRCh38 (3.04±0.33%, p=1.67×10^-13^) (**Fig. 2D**). It is obvious that YAO outperforms both CHM13 and GRCh38 in WES data analysis when Chinese samples are mapped, even in the case where the capture probes are not appropriate for Chinese samples.

### Improvement in germline variant calling

Using DNAscope, an accurate and efficient germline small-variant caller that combines the mathematics of the GATK’s HaplotypeCaller with a machine-learned genotyping model [28], we called germline variants. Generally, homozygous variants have a frequency close to 1, heterozygous variants around 0.5, while somatic variants exhibit frequencies deviating from 0.5 and 1. Based on these general rules, variants are further determined using deep learning models that consider additional factors such as depth, base quality, and mapping quality. The raw variant results are filtered by default cutoffs of >30×depth and >30 quality score to generate a list of high-confidence variants (see flowchart). The number of germline variants decreases significantly when using YAO as a reference compared to the other two references, for both homozygous and heterozygous variants (**Fig. 3AB**). Since homozygous variants often have high frequency in population, and thus are most likely associated with population-specific variations, we only identified 715,828±149,696 such variants when using YAO as a reference. However, this number increased by 11.95% and 19.26% when CHM13 (801,369±119,952; p=3.58×10^-15^, paired t-test) and GRCh38 (853,687±161,424; p=3.55×10^-17^, paired t-test) are used as references, respectively (**Fig. 3A**). For heterozygous variants, which are primarily attributable to within-population diversity and low-frequency variations, we identified similar number of variants when referring to YAO and CHM13, which are 729,123 ± 191,013 and 735,117 ± 152,423, respectively. However, GRCh38 still ensures an identification of heterozygous variants as many as 777,471±200,933, which is 6.62% more than YAO does (*p*=3.65×10^-13^, paired t-test, **Fig. 3B**). Compared to GRch38, more variants were shared using YAO and CHM13 as reference genomes (**Fig S3**). After further filtering out varinats with low-quality score (<30) or those in regions with lower reads depth (< 30×), homozygous germline variants obtained using YAO are also the fewest among the three reference genomes (**Fig S4A**). However, the differences in count of heterozygous variants obtained from the three reference genomes were reduced (**Fig S4B**).

**Fig 3.**
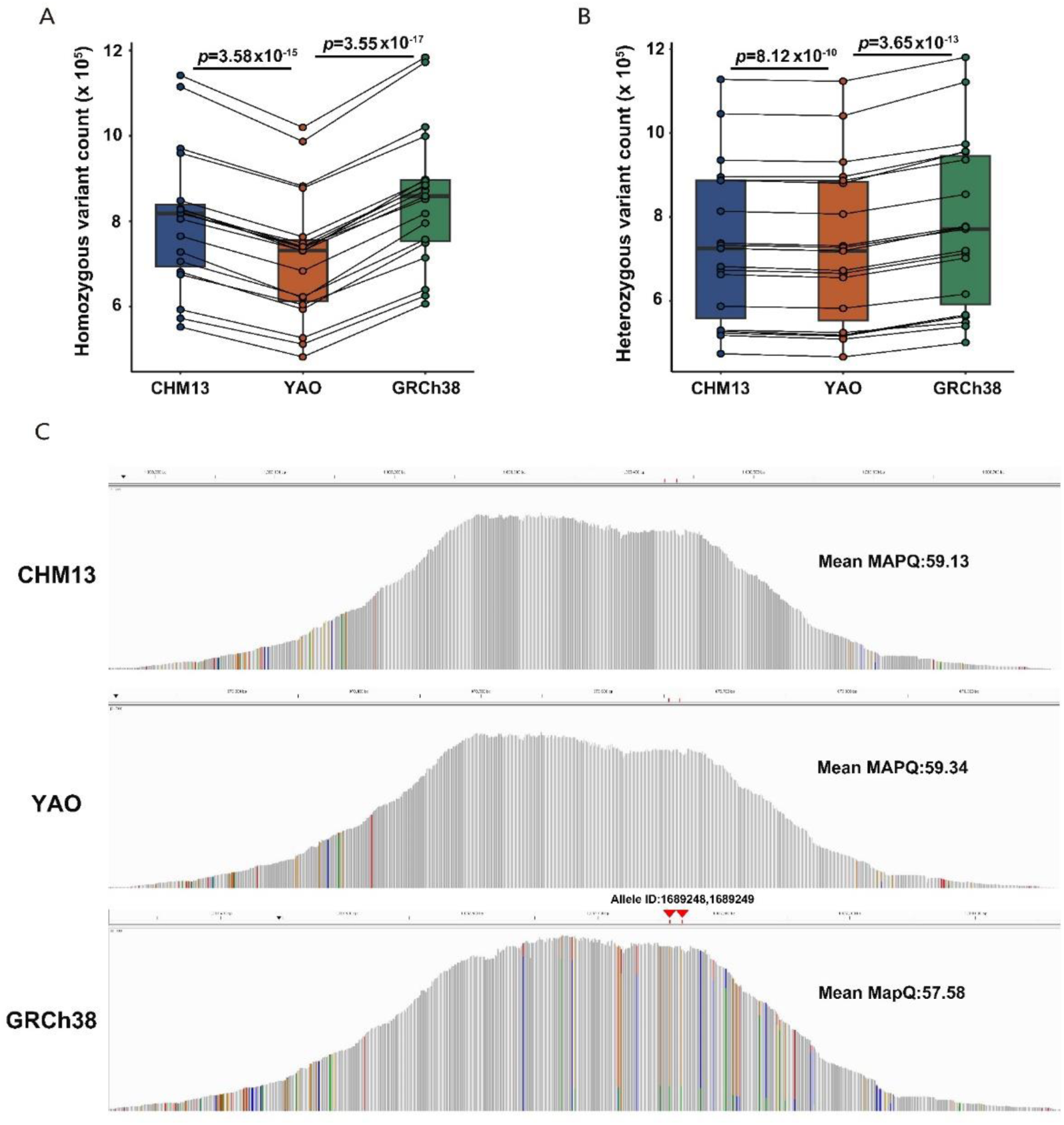
Comparison of germline variants referencing CHM13, YAO, and GRCh38. **A.** Total homozygous variants called by DNAscope in all reads covered regions. **B.** Total heterozygous variants called by DNAscope in all reads covered regions. The *p* value of paired T-test is labeled above each comparison. The points representing the same individual across different reference genome are connected by solid lines. **C.** Peak plot of reads mapped to the target region of 7^th^ exon in *CNN2* gene. The horizontal coordinate of each vertical line represents the position of each base, and the length of the vertical line represents the sequencing depth. Gray lines indicate that the bases are consistent with those in the reference genome, and colored lines indicate inconsistencies with bases in the reference genome. The colors green, red, orange, and blue represent the four bases, A, T, G, and C, respectively. Red arrows indicate pathogenic variants of allele ID 1689248 and 1689249 (recorded in the ClinVar) in GRCh38 and there are no variants on its corresponding sites in YAO and CHM13.

Similarly, when narrowing down to the probe targeted region, we found the same trends. There are the fewest homozygous variants called using YAO as references (**Fig.S5**). Comparing YAO and CHM13, the findings indicate that population-associated variations are a primary factor contributing to the identification of homozygous variants in samples. The difference between YAO and GRCh38, a chimeric genome, is slightly wider, possibly due to the more assembly errors in GRCh38 that are not present in any population and thus lead to more homozygous variant calls.

Further scrutinizing the different variants calls using the three references, we observed no YAO-specific or CHM13-specific pathogenic variants reported in ClinVar database (v.20231121, see below) [29] but identified four GRCh38-specific pathogenic ones, with two located in the 7th exon of the *CNN2* gene transcript NM_004368.7 (encoding calponin 2) on chromosome 19, found in 13 out of 19 samples. We further examined reads mapped to this exon from one sample (sample St-2) harboring the two variants, and found that using YAO and CHM13 as reference, the reads were well-mapped with few mismatches. However, using GRCh38 as reference, an additional subset of reads was mapped to this region, bearing numerous mismatches. Tracing these additional reads using YAO and CHM13 as reference, they were primarily from a pseudogene located in the pericentromere region of chromosome 20 **(Fig S6AB**), which is buried under many tandem repeats. This pseudogene is partially homologous to the exon of CNN2 and is absent in GRCh38 due to the poorly assembled pericentromere region in chromosome 20. As a result, when using GRCh38 as the reference, reads from this pseudogene were misaligned to the CNN2 exon, leading to many false positives, including the pathogenic variants (**Fig 3C, Fig S6C**). It illustrates how structural variations between reference genomes can affect read mapping and result in false-positive variant calls.

### Assessment in identifying pathogenic variants

For better interpreting clinically-significant variants, we use ANNOVAR to screen the records in the ClinVar database (v.20231121) [29], containing a total of 2,336,658 records. When converting the ClinVar coordinates from GRCh38 to CHM13 and YAO, only 5,186 (0.22%) and 5,967 (0.26%) records failed conversion for YAO and CHM13, respectively. However, we have only hit numbers per sample 14407.9 ± 725.5 for YAO and 16618.5 ± 834.5 for CHM13, in contrast to a much higher hitting rate for GRCh38, 31526.7 ± 1542.9 per sample (**Table S4**). This added difference is largely attributed to the categories of *Benign*, *Likely benign*, and *Conflicting interpretations of pathogenicity* (**Fig. 4A**). Furthermore, a much larger proportion of the variants in *Benign* categories are homozygous in GRCh38 (48.01%), compared to YAO (28.92%) and CHM13 (31.24%). The reduced counts in YAO and CHM13 are mostly from homozygous variants found in the categories of *Benign* and *Likely benign*. The differences observed in the number of variants annotated as related to clinical phenotypes by ClinVar between YAO and GRCh38 may be attributed to several factors, including population-specific variants, particularly those that are homozygous or with high frequency, as well as false positives, as illustrated in Fig. 3C. ClinVar annotations are based on GRCh38, which is a mosaic reference genome created by merging data from multiple donors. This approach generates an excess of artificial haplotypes and rare alleles, potentially introducing subtle biases in the analysis [19]. Consequently, using GRCh38 as a reference may result in a higher number of homozygous variant calls. Additionally, assembly errors or copy number variations in GRCh38 might contribute to false positive calls, leading to an increased number of variant annotations, including a higher frequency of benign records.

**Fig 4.**
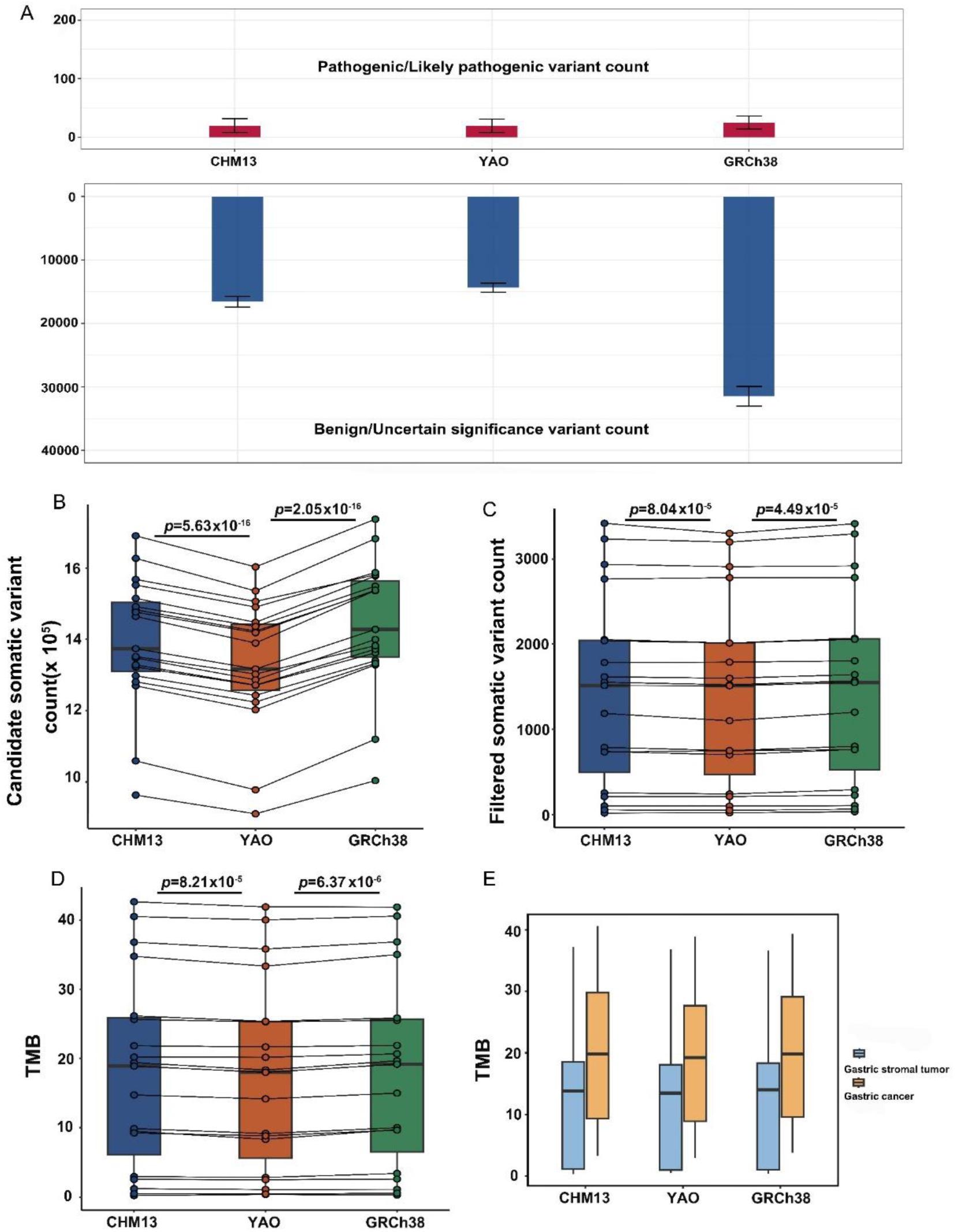
Comparison of clinically relevant variants and tumor mutation burden when using CHM13, YAO, and GRCh38 as references. **A.** Count of average ClinVar recorded *Pathogenic* (upper panel) variants and the sum count of *Benign* and *Uncertain significance* (lower panel) variants. **B.** Count of candidate somatic variants detected by TNScope. **C.** Count of somatic variant after filter. **D.** Tumor Mutational Burden (TMB). The *p* value of paired T-test is labeled above each comparison. The points representing the same individual across different reference genome are connected by solid lines. **E.** TMB comparison between gastric stromal tumors and cancers.

YAO, when applied to Chinese population samples, achieves similar sensitivity in identifying pathogenic variants in addition to reducing false positives. In the categories of *Pathogenic* and *Likely pathogenic*, we identified similar numbers of variants using either YAO (19.4 ± 11.5 per sample) or CHM13 (19.9 ± 11.6 per sample), a little less than that when using GRCh38 (25.2 ± 10.9 per sample). Upon scrutinizing the *Pathogenic* variations, we found that many variations in GRCh38 are attributable to reads with wrong mapping, whereas no such variants are seen in the counterpart positions in YAO (**Fig 3C**). This discrepancy may arise from incorrect mapping in GRCh38, whilst they are accurately recognized in YAO.

### Evaluation in TMB analysis

We further analyzed somatic variants and calculated TMB, which is composed of a standardized number of non-synonymous mutations and serves as an indicator for the presence of tumor-specific antigens, capable of predicting treatment responses in cancer immunotherapies [30]. For calculating TMB, blood samples are typically required to exclude germline variants from those identified in tumor tissues, thereby reducing false-positive calls of somatic variants. However, as many of the FFPE samples we utilized did not have corresponding blood samples available, we employed only the tumor WES data for somatic variant calling. We first utilized the tumor-only mode TNscope [31] to call candidate somatic variants in target exon regions, and subsequently removed those also identified as germline variants by DNAscope. The somatic variant sets were further filtered with read depth (>30× within the target exon region), quality score (>30), and a false-positive-specific filter tool FPfilter [32]. We picked up the non-synonymous mutations according to annotations from the filtered somatic variants and calculated TMB standardized by the length of WES.

As expected, TNscope identified significantly fewer candidate somatic variants when YAO was used as a reference as compared to the other two references (**Fig. 4B**). However, the difference is substantially reduced after removing germline variants and the application of other filters, yet it remains significant (**Fig. 4C**). The final YAO-based TMB values are slightly but significantly lower than those based on the other two references, possibly due to fewer false-positive somatic variant calls (**Fig. 4D**). For all reference genomes, samples of malignant gastric cancer exhibit higher TMB values than those of benign stromal tumor, indicating higher TMB for malignant cancers, though this difference was not statistically significant due to the small sample size (**Fig. 4E**). And due to the absence of normal samples and consequent inadequate germline variation filtering, the TMB of tumor-only WES is slightly higher than previously reported in gastric cancer and gastric stromal tumors [33, 34].

### Limitations of this study

Limitations of this study include the use of FFPE samples, which introduce variability in tumor purity and data quality, and a small sample size that restricts the statistical significance of our analysis of disease-related variants and their clinical relevance between benign gastric stromal tumors and malignant gastric cancers. Additionally, the absence of normal samples may have led to incomplete removal of germline variants, resulting in a slightly elevated tumor mutational burden (TMB), despite our rigorous filtering methods. Nevertheless, the primary goal of this study was to preliminarily assess the performance of different reference genomes, focusing on the utility of population-specific references in the upcoming T2T era. The findings reveal significant differences when using alternative reference genomes compared to GRCh38, underscoring the need for further optimization of variant calling processes and the accumulation of genomic data from the Chinese population. Such advancements will improve the identification of disease-related variants and enhance the clinical applicability of indices like TMB.

### Conclusions

This study conducts a parallel comparison of the current human reference genome GRCh38 and potential reference genomes with top-quality—YAO and CHM13—in the whole process of genomic analysis of WES data from 19 tumor samples of Chinese patients using state-of-the-art algorithms and tools. The initial comparison reveals that the three reference genomes share similar basic characteristics in terms of genome size, GC content, and exome proportion, except GRCh38 which is incomplete with many unfilled gaps, and its mosaic nature leads to inevitable misassembled contigs. Subsequent analyses of WES data illustrated that both YAO and CHM13 outperform GRCh38 as a reference by offering higher mapping rates, lower mismatch rates, and more reliable variant calling and annotation. Our initial study demonstrates that YAO, with quality similar to CHM13, is more suitable for samples from Chinese individuals, thus proposing the idea of the population-specific reference genome. The reads mapping results demonstrate the effectiveness of YAO in accurately aligning sequencing reads to the reference genome, ensuring high-quality data for downstream functional analysis. The high mapping rate and coverage of YAO as a reference genome for population-based studies for Chinese patients underscore its suitability, especially in clinical settings and for disease treatments.

## Materials and Methods

### Data collection and alignment

In this study, quality control of the sequencing data was performed using FastQC v0.11.8 (https://github.com/s-andrews/FastQC) to assess the quality of raw sequencing reads and identify potential issues, and MultiQC [35] was employed to generate a comprehensive report. To ensure the complete removal of adapter sequences and low-quality bases, sequencing reads were processed using TrimGalore-0.6.10 (https://github.com/FelixKrueger/TrimGalore). Sample alignments to the reference genomes, CHM13v2.0, YAO, and GRCh38.p14, were performed using BWA-MEM v0.7.17-r1188 (https://github.com/lh3/bwa). After alignment, the sorting and PCR duplicate removal of BAM files were processed with SortSam and MarkDuplicates commands of Picard tools v3.1.0 (https://github.com/broadinstitute/picard).

### Alignment quality assessment

To analyze the alignment results, we first used the stats command of samtools tool (version 1.9) [36] to extract various alignment parameters. Next, we used samtools depth command to extract the coverage and depth of the alignment results across the whole genome. In addition, we utilized transanno (https://github.com/informationsea/transanno) to lift the coordinates of the exome probe regions from GRCh37 to the other three reference genomes. This step is essential for comparing the alignment results and analyzing the exome region across different reference genomes. To compare exon regions, the corresponding exon probe sequences were aligned with the Needle tool within the EMBOSS suite [37] to determine the percentage identity between the sequences.

### Variant calling and variant annotation

DNAScope [28] was used to identify germline variants. Variants rejected by the machine learning algorithms in DNAScope were filtered out. For further filtering, germline variants with a quality score below 30 or depth below 30 are removed. The tumor-only mode in TNScope was utilized to identify somatic variants [31]. Variants that failed to pass the criteria mentioned above or shared by DNAScope variant calling were removed, and a final filtering step was performed using FP-filter to identify somatic variants. The ClinVar_20231126 database [29] was downloaded, and databases specific to CHM13 and YAO were established using transanno and the Vt toolkit [38]. The ANNOVAR tool [39] was used for variant annotation.

## Ethical statement

The application for the study was submitted to and approved by the Ethical Review Committee of Linfen Central Hospital (Approval No. YP2023-47-1). The collection and storage of human samples were registered with and approved by the Human Genetic Resources Administration of China (HGRAC). Written informed consents were obtained from all participants.

## Code availability

The code of this work is available on GitHub (https://github.com/KANGYUlab/WES).

## Data availability

The raw WES data of 19 fresh gastric tumor samples have been deposited in the GS A for human in China National Center for Bioinformation (CNCB) under the accession number HRA006227 which is publicly accessible at https://ngdc.cncb.ac.cn/gsa-human. The T2T-YAO.hp genome is available at NGDC Genome Warehouse (https://ngdc.cn cb.ac.cn/gwh/) (GWHDQZI00000000). T2T-CHM13v2.0 is available at NCBI (GCA_00 9914755.4). GRCh38 genome and its annotation file are available at UCSC (GCA_000 001405.15, https://hgdownload.soe.ucsc.edu/goldenPath/hg38/bigZips/hg38.fa.gz; https://hgdownload.soe.ucsc.edu/goldenPath/hg38/bigZips/genes/hg38.refGene.gtf.gz). The VCF file s containing filtered variants of each sample called by DNAScope and TNScope in thi s paper are available on GitHub (https://github.com/KANGYUlab/WES)

## CRediT authorship contribution statement

<u>Shuming Guo</u>: Investigation, Resources, Data curation, Formal analysis, Funding acquisition, Writing – original draft. <u>Zhuo Huang</u>: Investigation, Methodology, Data curation, Formal analysis, Software, Visualization, Writing – original draft, Writing – review & editing. <u>Yanming</u> <u>Zhang</u>: Investigation, Resources, Data curation, Formal analysis, Writing – original draft. <u>Yukun He</u>: Resources, Data curation, Formal analysis, Writing – original draft. <u>Xiangju Chen:</u> Resources, Writing – original draft. <u>Wenjuan Wang</u>: Resources, Writing – original draft. <u>Lansheng Li:</u> Resources, Writing – original draft. <u>Yu Kang</u>: Conceptualization, Formal analysis, Investigation, Methodology, Project administration, Supervision, Writing – original draft, Writing – review & editing. <u>Zhancheng Gao</u>: Funding acquisition, Project administration, Supervision, Writing – review & editing. <u>Jun Yu</u>: Conceptualization, Formal analysis, Funding acquisition, Investigation, Methodology. <u>Zhenglin Du</u>: Investigation, Methodology, Data curation, Formal analysis, Writing – original draft, Writing – review & editing. <u>Yanan Chu</u>: Conceptualization, Funding acquisition, Investigation, Methodology, Data curation, Formal analysis, Writing – original draft, Writing – review & editing.

## Acknowledgments

This study was supported by the National Key Research and Development Program of China (Grant No. 2021YFC2301000), the National Science Foundation of China (Grant No. 32371537), and the grants of Linfen Soft Science Research Project (Grant No. 2126). National and Provincial Key Clinical Specialty Capacity Building Project 2020 (Department of the Respiratory Medicine), and Peking University People’s Hospital Scientific Research Development Funds (Grant No. RDGS2022-11)

## Supplementary material

**Fig S1. Analysis pipeline.**

**Fig S2. Comparison of all mapped reads alignment between YAO vs. GRCh38 and YAO vs. CHM13.**

**A.** The reduction of all aligned reads referencing CHM13 and GRCh38 compared to YAO in whole genome region. Blue and orange solid circles represent differences between YAO vs GRCh38 and YAO vs CHM13, respectively. Samples were sorted according to their total mapped reads. St, gastric stromal tumor; Ca, gastric cancer. **B.** Mismatch rates calculated as the number of mismatched bases divided by the total number of aligned bases. **C.** The comparison of coverage of each reference genome based on all aligned reads of each sample. The coverage was calculated as the length of reads covered regions with depth >= 1 divided by the total length of the reference genome. The p value of paired T-test is labeled above each comparison.

**Fig S3. Venn plots of germline variants across different reference genomes. A.** Unfiltered variants. **B.** Filtered variants.

**Fig S4. Comparison of filtered germline variants in all mapped regions referencing CHM13, YAO, and GRCh38. A.** Filtered germline homozygous variants. **B.** Filtered germline heterozygous variants. The p value of paired T-test is labeled above each comparison.

**Fig S5. Comparison of germline variants in target exon regions referencing CHM13, YAO, and GRCh38. A.** Total unfiltered germline homozygous variants in target regions. **B.** Total unfiltered germline heterozygous variants in target regions. **C.** Filtered germline homozygous variants in target regions. **D.** Filtered germline heterozygous variants in target regions. The p value of paired T-test is labeled above each comparison.

**Fig S6. The homologous regions and reads alignment of CNN2 gene exon No.7 in GRCh38, YAO, and CHM13. A**. The alignments between syntenic (chr19) and non-syntenic (chr20) homologous regions around CNN2 gene exon No.7 on GRCh38, YAO and CHM13. Homologous regions are linked by grey blocks between the chromosomes. **B**. Reads alignment in the non-syntenic homologous regions on chr20 in YAO and CHM13; **C**. Reads alignment in the region around CNN2 gene exon No.7 on chr19 in GRCh38. Pink or purple horizontal lines represent forward or reverse reads, respectively. Reads in the black dotted box are wrongly mapped to Chr19 in GRCh38, actually located on Chr20, around 30M in YAO and CHM13.

**Table S1. Comparison of basic statistics of CHM13, YAO, and GRCh38.**

**Table S2. Comparison of the WES target regions lifted from GRCh37 to CHM13, YAO, and GRCh38.**

**Table S3. Information of WES sequencing samples**

**Table S4. Comparison of the clinically relevant variants identified using CHM13, YAO, and GRCh38 as reference.**

